# Stress-Mediated Attenuation of Translation Undermines T Cell Tumor Control

**DOI:** 10.1101/2022.01.31.478547

**Authors:** Brian P. Riesenberg, Elizabeth G. Hunt, Megan D. Tennant, Katie E. Hurst, Alex M. Andrews, Lee R. Leddy, David M. Neskey, Elizabeth G. Hill, Guillermo O. Rangel Rivera, Chrystal M. Paulos, Peng Gao, Jessica E. Thaxton

## Abstract

Protein synthesis enables cell growth and survival, but the molecular mechanisms through which T cells suppress or maintain protein translation in the stress of solid tumors are unknown. Using mouse models and human tumors we demonstrate that protein translation in T cells is repressed by the solid tumor microenvironment (TME) due to activation of the unfolded protein response (UPR) via phosphorylation of the α subunit of eukaryotic translation initiation factor 2 (p-eIF2α). Given that acute glucose deprivation in T cells exacerbated p-eIF2α, we show that metabolic reprogramming toward glycolytic independence allays the UPR and p-eIF2α, enabling sustained protein translation in T cells in TME stress. UPR mitigation was associated with enhanced degradation of proteins in antitumor T cells, as proteasome inhibition resulted in eIF2α phosphorylation, attenuation of translation, and loss of antitumor efficacy. In contrast, proteasome stimulation relieved translation inhibition, inducing robust T cell tumor control, offering a new therapeutic avenue to fuel the efficacy of tumor immunotherapy.

## Introduction

Activation of protein synthesis is a requirement for T cell growth and effector function [1]. Eukaryotic translation initiation factor 2 (eIF2) controls cap-dependent protein translation efficiency by bridging Met-tRNAi and the ribosomal subunit [2]. However, endoplasmic reticulum (ER) stress, catalyzed by accrual of unfolded proteins in the ER lumen, undermines the competency of the process. In response to ER stress, the unfolded protein response (UPR) is initiated via phosphorylation of the α subunit of eIF2 causing translation attenuation as a means to restore proteostasis [3]. The tumor microenvironment (TME) is replete with metabolic stressors known to activate the UPR [4-7]. Indeed, we and others have shown that PKR-ER-like kinase (PERK), a stress sensor responsible for eIF2α phosphorylation [8, 9], undermines T cell antitumor efficacy [10, 11]. While these studies implicate the UPR-mediated translational machinery as a potential molecular checkpoint prompted by TME stress, the extent to which translational regulation influences outcomes in the context of antitumor immunity is unknown.

Glycolysis is the critical energy requirement for T cells to undergo protein translation [12]. However, upon entering the TME CD8 T cells encounter competition for exogenous glucose resulting in a significant reduction in effector function [5]. In contrast, T cells that depend on metabolic pathways apt for cell survival in nutrient stress, such as gluconeogenesis [13] or fatty acid oxidation [14], demonstrate heightened tumor control [15, 16]. Metabolic reprogramming through cytokine conditioning [17] or chronic glucose deprivation [15] generates T cells enriched for such pathways that fuel sustenance in nutrient deplete settings. We previously ‘demonstrated that metabolic reprogramming through cytokine conditioning generates T cells adept to sustain protein translation in solid tumors [18]. However, the mechanism by which translation is maintained in tumor stress is undefined.

The proteasome is a proteolytic complex responsible for the degradation of ubiquitinated proteins [19]. Proteasome inhibition exacerbates ER stress and promotes the UPR, rendering tumor cells susceptible to apoptosis [20]. The relationship between protein translation and degradation is symbiotic, as effective protein catabolism precludes activation of the UPR. Memory T cells, the T cell subset with heightened antitumor efficacy, are enriched for proteasome subunits and exhibit accelerated protein degradation [21]. Activation of the proteasome promotes memory T cell lineage development, linking protein degradation to memory fate [22]. Indeed, memory like T cells are capable of sustaining protein synthesis under TME stress; however, the synergy between sustained translation, protein degradation, mitigating the UPR, and optimal antitumor immunity has not been investigated.

In the present study we investigated the contribution of stress-mediated attenuation of translation to facilitate inhibition of efficacious antitumor T cells. We found that the TME activates the UPR via p-eIF2α to restrict protein synthesis and tumor control in T cells. Metabolic stress, fueled by glucose deprivation, was alleviated by metabolic reprogramming that mitigated the UPR, promoting continued translation in the TME. In TME stress, we demonstrate that antitumor translation was complimented by robust proteasome activation that protected T cells from signaling the UPR and subsequent translation attenuation, proving critical for tumor control. The data indicate that protein degradation underlies antitumor metabolism and translation that can be harnessed to amplify T cell tumor control.

## Results

### Protein Synthesis is attenuated in tumor infiltrating T cells

Recent advances in single cell RNA [23] and ATAC sequencing [24] have allowed for identification of genetic and epigenetic traits associated with enhanced antitumor T cell function. However, the mechanisms responsible for controlling the translation of these instructions into effector functions have yet to be elucidated in antitumor immunity. We recently published an assay that allows monitoring protein synthesis on a per-cell basis [18]. The fluorescent analogue of methionine, L-homopropargylglycine (L-HPG), is incorporated into new forming polypeptide chains and quantified by flow cytometry through Click-IT chemistry [1, 25-27] (**Fig 1a**). Using this approach, we assessed global protein synthesis in endogenous CD8 T cells across multiple organs in B16-F1 melanoma-bearing mice. Rates of translation were determined by normalizing to samples treated with translation inhibitor cycloheximide (CHX). Compared to splenic and tumor-draining lymph nodes (tDLN), tumor infiltrating CD8 T cells demonstrated a significant reduction in protein translation (**Fig 1b**). We replicated this phenomenon using freshly isolated human tumors from various cancer types and patient matched PBMCs suggesting that blunted protein synthesis is a common theme of tumor infiltrating CD8 T cells in both humans and mice (**Fig 1c**).

**Figure 1.**
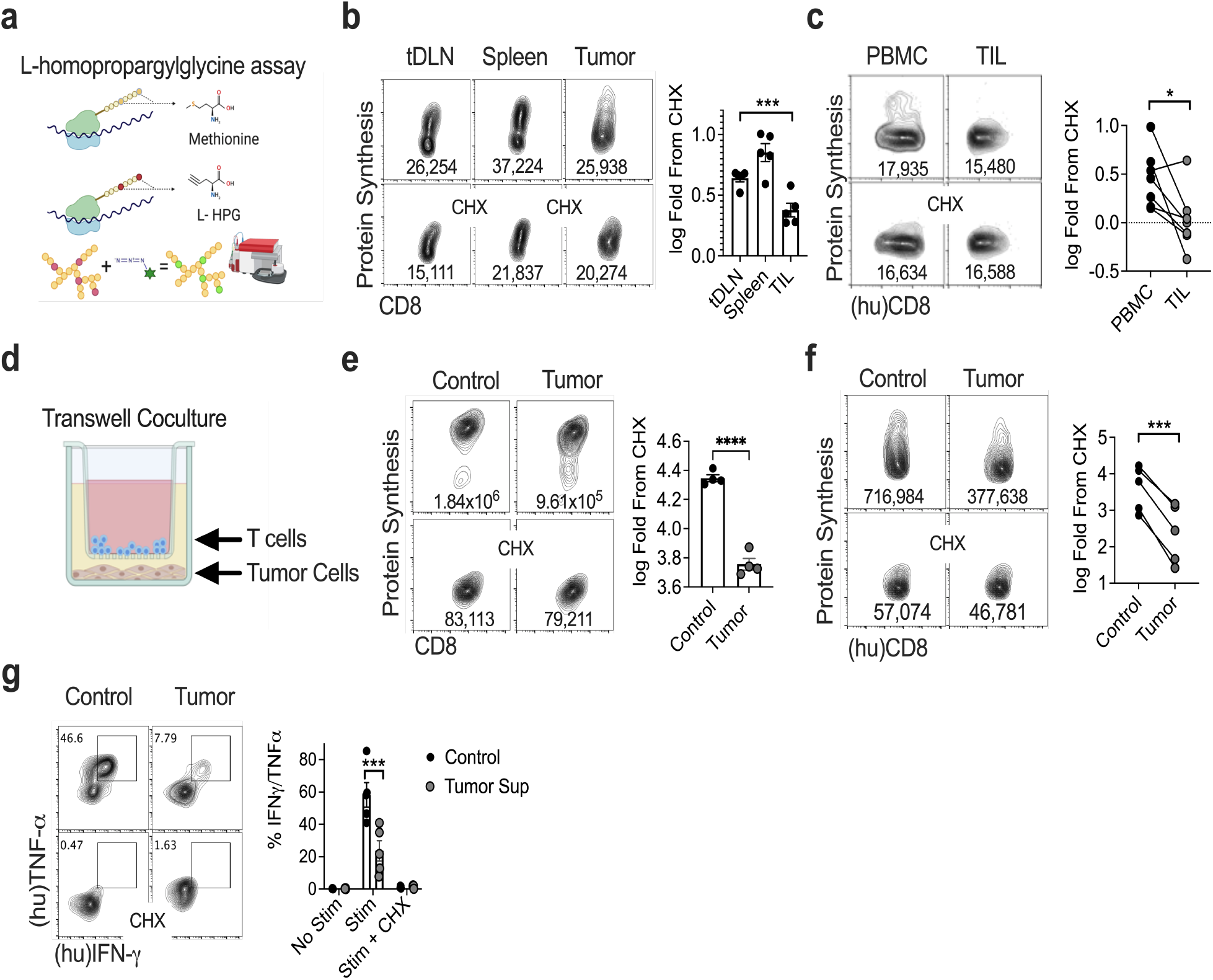
Protein Synthesis is attenuated in tumor infiltrating T cells. **a**) Schematic representation of L-homopropargylglycine assay (protein synthesis assay). **b**) Representative FACS plots and quantification of protein synthesis rates in b) endogenous CD8+ tDLNs, splenocytes, and TILs from B16F1 tumor-bearing mice (n=5 mice). **c**) Representative FACS plots and quantification of protein synthesis rates from CD8+ TILs and autologous PBMC from n=7 cancer patients. **d**) Graphic of tumor-T cell contact independent transwell coculture assay. Representative FACS plots and quantification of protein synthesis rates from **e**) freshly isolated OT-1 splenocytes stimulated with OVA peptide or **f**) CD8+ PBMC activated with CD3/28 activators as (n=5 donors) in the tumor transwell coculture. **g**) Representative FACS plots and quantification of IFN*γ* and TNFα production from CD8+ PBMC activated with CD3/28 activators (n=5 donors) in the tumor transwell assay followed by PMA/Ionomycin stimulation ± pretreatment with protein synthesis inhibitor cycloheximide (CHX). Data represent three to five independent experiments. *p < 0.05, **p < 0.01, ***p < 0.001, ****p < 0.0001 by one-way ANOVA (**b**), unpaired Student’s t-test (**e**) or paired Student’s t-test (**c, f, g**). (hu) indicates human samples, Error bars indicate the S.E.M.

The TME consists of a heterogenous milieu of cell types and biochemical processes that coordinate to suppress T cells in both contact independent and dependent manners [28, 29]. To resolve the critical aspects restricting protein translation in the TME, we utilized a coculture assay in which tumor cells were seeded prior to introducing T cells into transwell inserts [18] (**Fig 1d**). In tumor-seeded transwells, OT-1 T cells (**Fig 1e**) or normal human donor PBMC (**Fig 1f**) were activated and expanded with cognate OVA antigen or soluble CD3/28 activations, respectively, and protein translation rates were measured. In both mouse and human T cells the presence of tumor significantly reduced protein translation rates (**Fig 1e-f**). This phenomenon was also apparent when human CD8 T cells were expanded in the presence of supernatant from freshly isolated B16 tumors (**Supplemental Fig 1**). We further validated these observations using a second protein translation assay that incorporates O-propargyl-puromycin into the ribosomal A-site allowing for fluorescent labelling of nascent polypeptide chains (**Supplemental Fig 2**). These data indicate that tumor mediated reduction in protein translation occurs in part through a contact independent mechanism.

**Figure 2.**
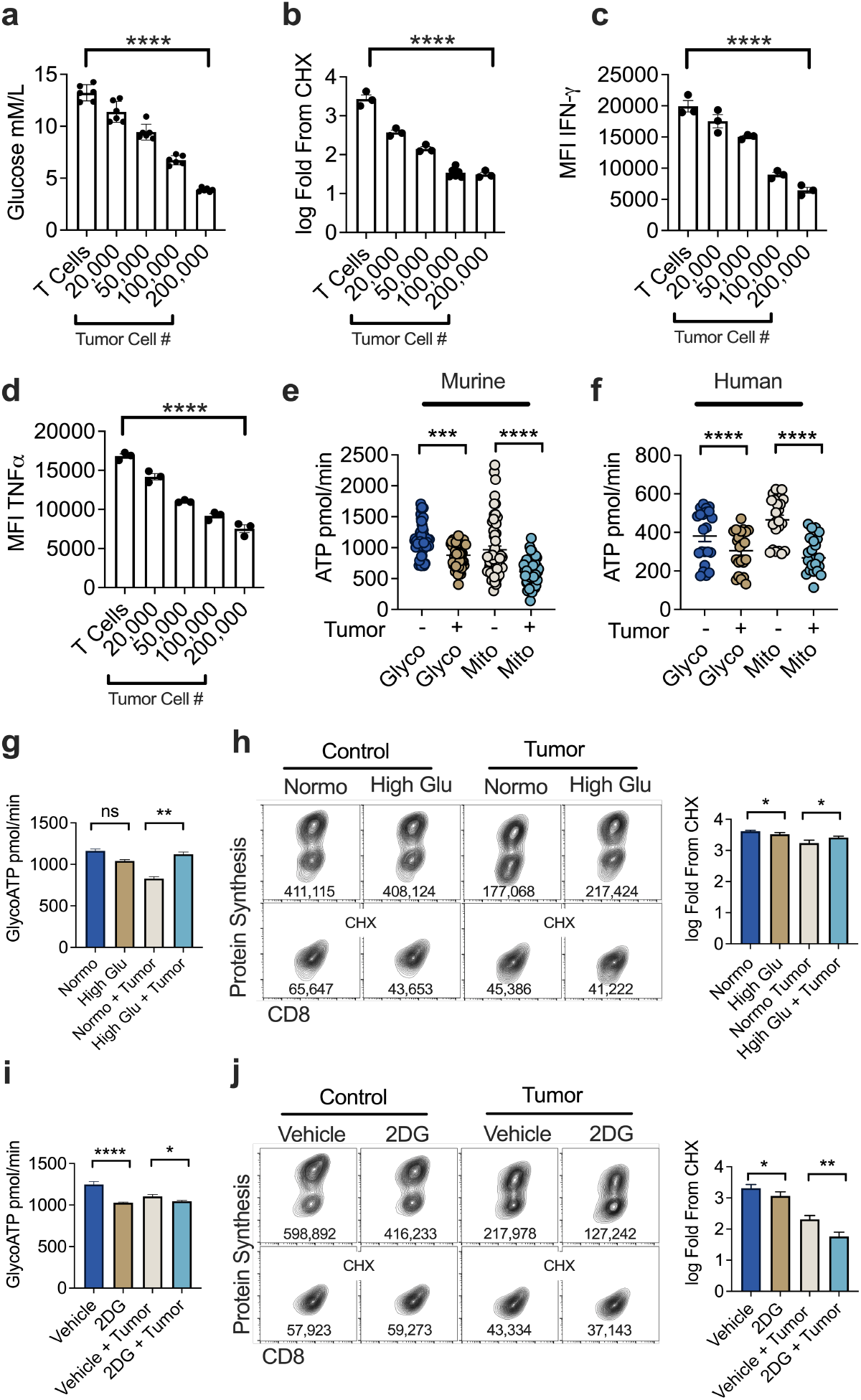
Glucose stress undermines T cell translation. Quantification of **a**) exogenous glucose concentration, **b**) protein synthesis rates, **c**) IFN*γ* or **d**) TNFα production from the tumor transwell assay with an increasing number of tumor cells seeded 24 hours prior to introduction of T cells. ATP production from glycolysis (glycoATP) or mitochondria (mitoATP) from **e**) OT-1 murine or **f**) human CD8^+^ PBMC (n = 4 donor) in the tumor transwell assay. Quantification of **g**) ATP production from glycolysis or **h**) protein synthesis rates with or without the addition of exogenous glucose (25mM) from the tumor transwell assay. Quantification of **i**) ATP production from glycolysis or **j**) protein synthesis rates with 2-deoxy-glucose (2DG) or vehicle added 2 hours prior to harvest in the tumor transwell assay. Unless otherwise specified, glucose measurements, ATP rate data, and protein synthesis measurements represent one of three to six independent experiments. *p < 0.05, **p < 0.01, ***p < 0.001, ****p < 0.0001 by one-way ANOVA (**a-d**), unpaired (**e, g-j**) or paired Student’s t-test (**f**). Error bars indicate the S.E.M.

Given that cytokine synthesis is a product of protein translation [30] we asked whether the reduction in protein translation in the TME corresponded with diminished cytotoxic cytokine synthesis. Upon coculture with tumor cells, human PBMCs were activated with soluble CD3/28 followed by cytokine restimulation. Coculture with tumor cells diminished TNFα/IFN*γ* production in CD8 PBMCs (**Fig 1g**). Pretreatment of cells with CHX prior to restimulation resulted in the complete abrogation of TNFα/IFN*γ* in control and TME CD8 PBMCs, indicating the requirement of continuous translation for cytokine synthesis. Cytokine production in the presence of tumor cells was also reduced using the OT-1 system to validate our findings (**Supplemental Fig 3**). Together, our data illustrate that contact independent TME stress undermines cytokine synthesis in CD8 T cells.

**Figure 3.**
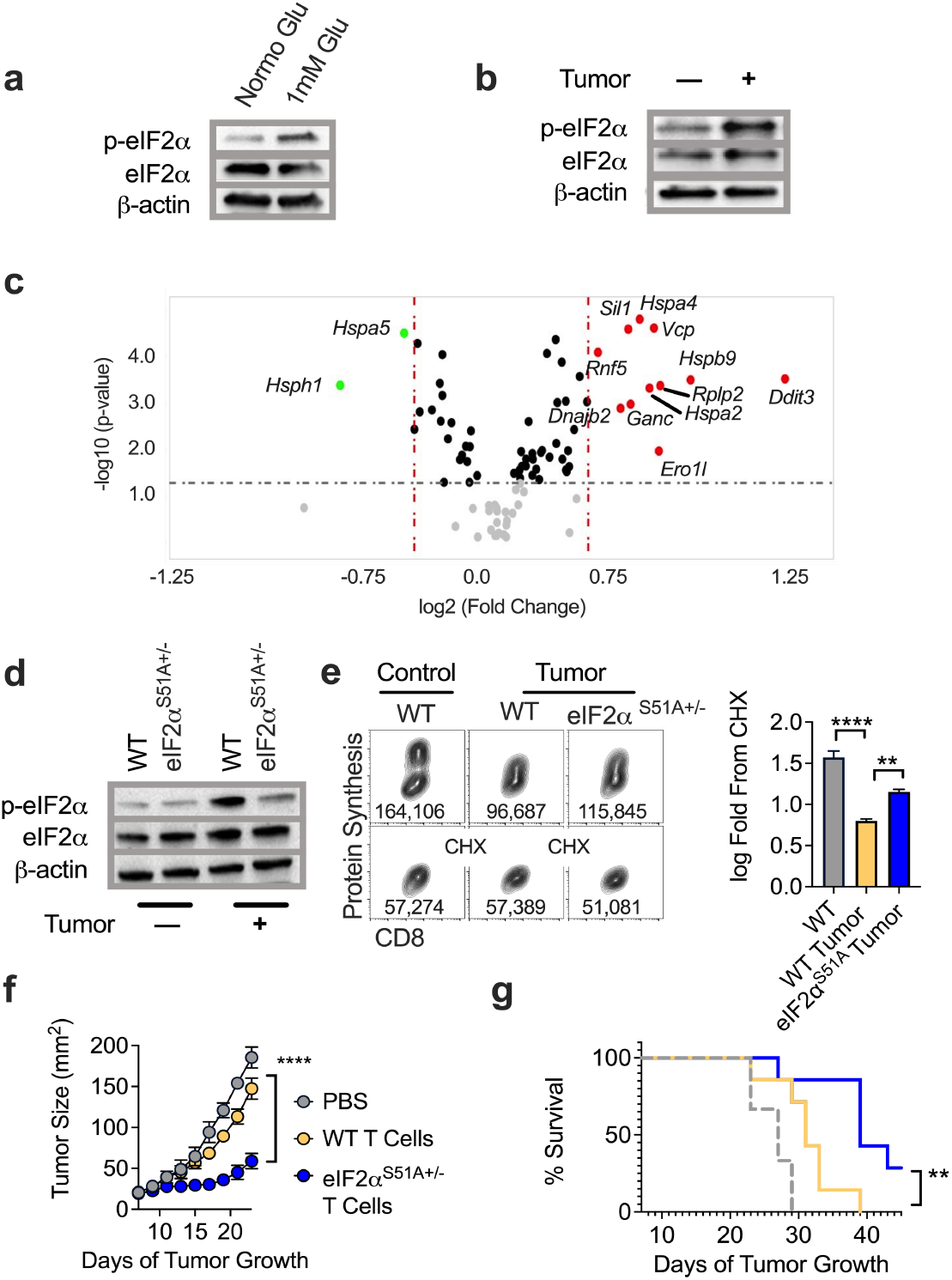
eIF2α phosphorylation attenuates T cell translation and tumor control. Western blot analysis of phosphorylated or total eIF2α with β-actin loading controls from OT-1 T cells cultured in **a**) normal or low glucose (1mM) conditions for 12 hours or **b**) the tumor transwell system with or without pre-seeded tumor cells. **c**) Volcano plot analysis from a targeted gene array measuring 83 common UPR associated genes from OT-1 T cells harvested from the tumor-T cell coculture assay with or without pre-seeded tumor cells. Genes in red are upregulated in the tumor condition while green are genes down regulated. **d**) Western blot analysis of phosphorylated or total eIF2α with β-actin loading controls from eIF2α^S51A+/-^ or wild type littermate OT-1 controls harvested from the tumor transwell assay. **e**) Representative FACS plots and quantification of protein synthesis rates from eIF2α^S51A+/-^ or wild type littermate OT-1 controls harvested from the tumor transwell assay. **f**) Tumor growth rate or **g**) overall survival from adoptive transfer of day 7 expanded eIF2α^S51A+/-^ or wild type littermate OT-1 T cells into day 7 B16-F1-OVA bearing B6 mice. Western blot and FACS plots represent one of three to six independent experiments. UPR gene array was performed with technical triplicates. Tumor control experiments represent one of two independent experiments. *p < 0.05, **p < 0.01, ***p < 0.001, ****p < 0.0001 by one-way ANOVA (**e**) or linear regression (**f**) and log-rank, mantel-cox test of survival proportions (**g**). Criteria for significance in the UPR gene array were Fold Change > 1.5 and p < 0.05. Error bars indicate the S.E.M.

### Glucose stress undermines T cell translation

Protein translation, particularly at the magnitude seen in effector T cell activation and expansion [21], is a metabolically intensive process requiring ATP [31]. A common stressor impacting T cell function in solid tumors is competition for exogenous glucose with tumors cells [5]. Given that exogenous glucose fuels effector cell metabolism, in addition to the contact independent nature of our previous findings, we asked whether competition could be driving the reduction in protein synthesis in T cells. Incremental increases in tumor cell numbers resulted in a stepwise reduction in environmental glucose in coculture media (**Fig 2a**). This was coupled with an almost identical graded reduction in global protein synthesis, IFN*γ*, and TNFα production (**Fig 2b-d**) highlighting the link between glucose availability and CD8 T cell functional capacity. Indeed, exposure to low glucose associated with TME pressure in the coculture system resulted in a significant reduction in ATP output, irrespective of the metabolic source, in both OT-1 mouse and purified human CD8 T cells (**Fig 2e-f**). Thus, competition for glucose significantly alters T cell metabolism, thereby limiting their effector function.

We performed a series of rescue or removal experiments to confirm the necessity of glucose in driving protein translation. First, by supplementing the T cell-tumor cell coculture media with exogenous glucose, we were able to restore glycolytic ATP output to levels similar to no tumor controls (**Fig 2g**). In line with this, exogenous glucose was also able to rescue protein translation rates in the presence of tumor pressure (**Fig 2h**). We next inhibited glycolysis in T cells in the transwell coculture system by adding 2-deoxy-glucose (2DG) 2 hours prior to measurement of protein translation. 2DG is taken up by glucose transporters [32] and competitively inhibits generation of glucose-6-phosphate in the second step of glycolysis. Seahorse bioanalysis indicated a reduction in glycoATP (**Fig 2i**) in all 2DG conditions irrespective of tumor cell pressure, validating acute treatment with 2DG blunts glycolysis. 2DG treatment significantly reduced translation rate in T cells compared to vehicle control in tumor and non-tumor settings (**Fig 2j**). Combined, these data highlight the importance of exogenous glucose in dictating metabolic programming, protein translation, and effector function.

### eIF2α phosphorylation attenuates T cell translation and tumor control

Environmental changes, particularly glucose deprivation, lead to disturbances in ER protein folding initiating activation of the UPR [33]. This process is driven in part by phosphorylation of eIF2α at Ser51 [34, 35], which reduces global translation in favor of maintaining proteostasis [36]. Indeed, OT-1 CD8 T cells exposed to low glucose conditions *in vitro* demonstrated a significant increase in p-eIF2α at Ser51 compared to normal glucose controls (**Fig 3a**). We noted a similar phosphorylation pattern when comparing CD8 T cells harvested from the tumor cell transwell condition to the non-tumor controls (**Fig 3b**). Finally, using a targeted gene array specific to UPR activation, we found T cells harvested from tumor-seeded transwells upregulated 11 of 83 common UPR genes cells relative to non-tumor controls (**Fig 3c**). These results indicate that glucose deprivation mediated initiation of the UPR may be the underlying mechanism restricting protein translation in the TME.

To determine whether p-eIF2α is the direct arbiter of translation attenuation in T cells exposed to the TME, we backcrossed TCR transgenic OT1 mice with mice bearing a single amino acid substitution of serine to alanine at codon 51 (S51A) in the phosphorylation site of the eIF2α protein [3]. Animals homozygous for S51A mutation die near birth due to hypoglycemia, thus we studied p-eIF2α regulation in mice bearing T cells heterozygous for the S51A mutation (eIF2α^S51A+/-^). Western blotting indicated a staunch reduction in p-eIF2α in eIF2α ^S51A+/-^ T cells exposed to TME stress (**Fig 3d**) which correlated with an increase in translational capacity relative to littermate controls (**Fig 3e**). Given that previous reports have demonstrated the ability sustain translation in TME stress propels T cell tumor control [18], we aimed to test the effect of diminished translation attenuation by p-eIF2α on antitumor immunity. Subcutaneous B16F1-OVA melanomas were established in C57BL/6 mice and WT or eIF2α^S51A+/-^ OT-1 T cells were infused to mice bearing 7-day established tumors. eIF2α^S51A+/-^ cells induced better tumor control compared to WT-matched control T cells resulting in a significant extension in animal survival (**Fig 3f-g**). Our findings suggest that elevated eIF2α phosphorylation, driven through glucose deprivation, reduces translational capacity and subsequent antitumor immunity of T cells.

### Metabolic reprogramming alleviates UPR-associated translation attenuation

An interesting paradigm exists whereby T cells metabolically reprogrammed away from exogenous glucose dependency demonstrate heightened tumor control marked by amplified cytotoxic cytokine production [15, 16]. We next asked whether relief of glucose stress through metabolic remodeling rewires the UPR as a means to sustain antitumor function in reprogrammed T cells. We administered acute (2 hour) or chronic (36 hour) 2DG to OT-1 T cells seeded in our tumor transwell coculture. Western blotting revealed a pronounced reduction in eIF2α phosphorylation at Ser51 in the chronic 2DG treatment compared to acute 2DG treatment or vehicle control (**Fig 4a**). In line with this, we noted that UPR genes activated in T cells by TME exposure (**Fig 3c**) were now downregulated by chronic 2DG remodeling, indicating that glucose stress arbitrated UPR activation (**Fig 4b**). Moreover, chronic 2DG remodeling prompted elevated translation in T cells experiencing TME driven nutrient deprivation (**Fig 4c-d**) relative to non-tumor controls. Seahorse bioanalysis indicated a profound increase in both glycolytic and mitochondrial ATP output in chronic 2DG remodeled T cells under tumor stress (**Fig 4e-f**), suggesting complete metabolic reprogramming that supported energy production in nutrient stress.

**Figure 4.**
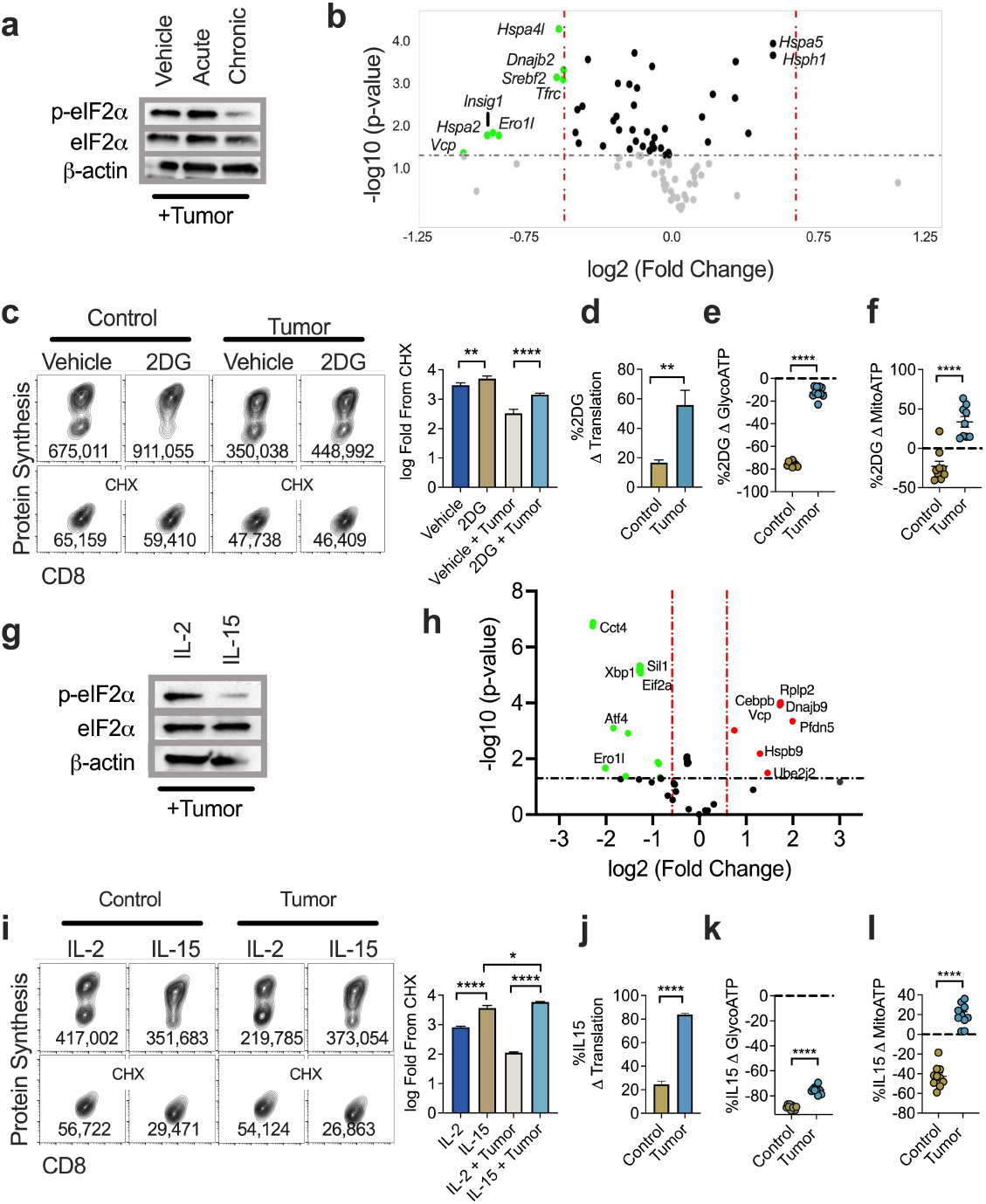
Metabolic reprogramming alleviates UPR-mediated translation attenuation. **a**) Western blot analysis of phosphorylated or total eIF2α with β-actin loading controls from OT-1 T cells cultured with vehicle, acute (2 hours), or chronic (36 hours) 2DG harvested from the transwell assay. **b**) Volcano plot analysis from a targeted gene array measuring 83 common UPR associated genes from OT-1 T cells harvested from the tumor-T cell coculture assay with chronic 2DG or vehicle control. Genes in green are down regulated in the chronic 2DG condition compared to vehicle control. **c**) Representative FACS plots and quantification of protein synthesis rates from chronic 2DG or vehicle treated controls harvested from the tumor transwell assay. Percent change in **d**) protein synthesis rates, **e**) glycolytic ATP or, **f**) mitochondrial ATP of chronic 2DG-treated OT1 T cells relative to vehicle control from tumor or non-tumor conditions in the tumor transwell assay. **g**) Western blot analysis of phosphorylated or total eIF2α with β-actin loading controls from OT-1 T conditioned with IL-2 or IL-15 prior to addition in the tumor transwell assay. **h**) Volcano plot analysis from a targeted gene array measuring 83 common UPR associated genes from OT-1 T cells conditioned with IL-2 or IL-15 prior to harvest from the tumor-T cell coculture assay. Genes in green are down regulated in the IL-15 condition relative to the IL-2 controls. **i**) Representative FACS plots and quantification of protein synthesis rates from IL-2 or IL-15 conditioned OT-1 T cells harvested from the tumor transwell coculture assay. Percent change in **j**) protein synthesis rates, **k**) glycolytic ATP or, **l**) mitochondrial ATP of IL-15 conditioned OT1 T cells relative to IL-2 control from tumor or non-tumor conditions in the tumor transwell assay. Western blot and FACS plots represent one of three to six independent experiments. *p < 0.05, **p < 0.01, ***p < 0.001, ****p < 0.0001 by two tailed Student’s *t* test (**c-f, i-l**). Criteria for significance in the UPR gene array were Fold Change > 1.5 and p < 0.05. Error bars indicate the S.E.M.

The possibility exists that T cell translation was bolstered by an adverse effect of chronic 2DG on tumor cells. Chronic 2DG conditioning promotes stem-like memory T cell development [16] which we confirmed using phenotypic analysis via flow cytometry (**Supplemental Fig 4)**. Thus, to validate elevated translation in TME stress was a property of metabolic remodeling of T cells to the memory lineage, we used the cytokine IL-15 to generate memory cells (**Supplemental Fig 5**). Comparable to chronic 2DG treatment, IL-15 memory-like T cells harvested from the tumor transwell coculture assay displayed a significant reduction in eIF2α phosphorylation when compared to IL-2 effectors (**Fig 4g**). Furthermore, we saw a similar reduction in the UPR when comparing IL-15 and IL-2 treated T cells exposed to the TME (**Fig 4h**), confirming IL-15 skewing alleviated UPR induced by TME pressure. IL-15 skewing enhanced protein translation rates in both non-tumor and tumor conditions, however, the effect was more pronounced in the nutrient deprived TME (**Fig 4i-j**). Moreover, IL-15 treatment produced a robust increase in ATP production in the presence of tumor pressure (**Fig 4k-l**). Taken together, these data definitively demonstrate that metabolic reprogramming mitigates UPR-mediated translation attenuation in T cells.

**Figure 5.**
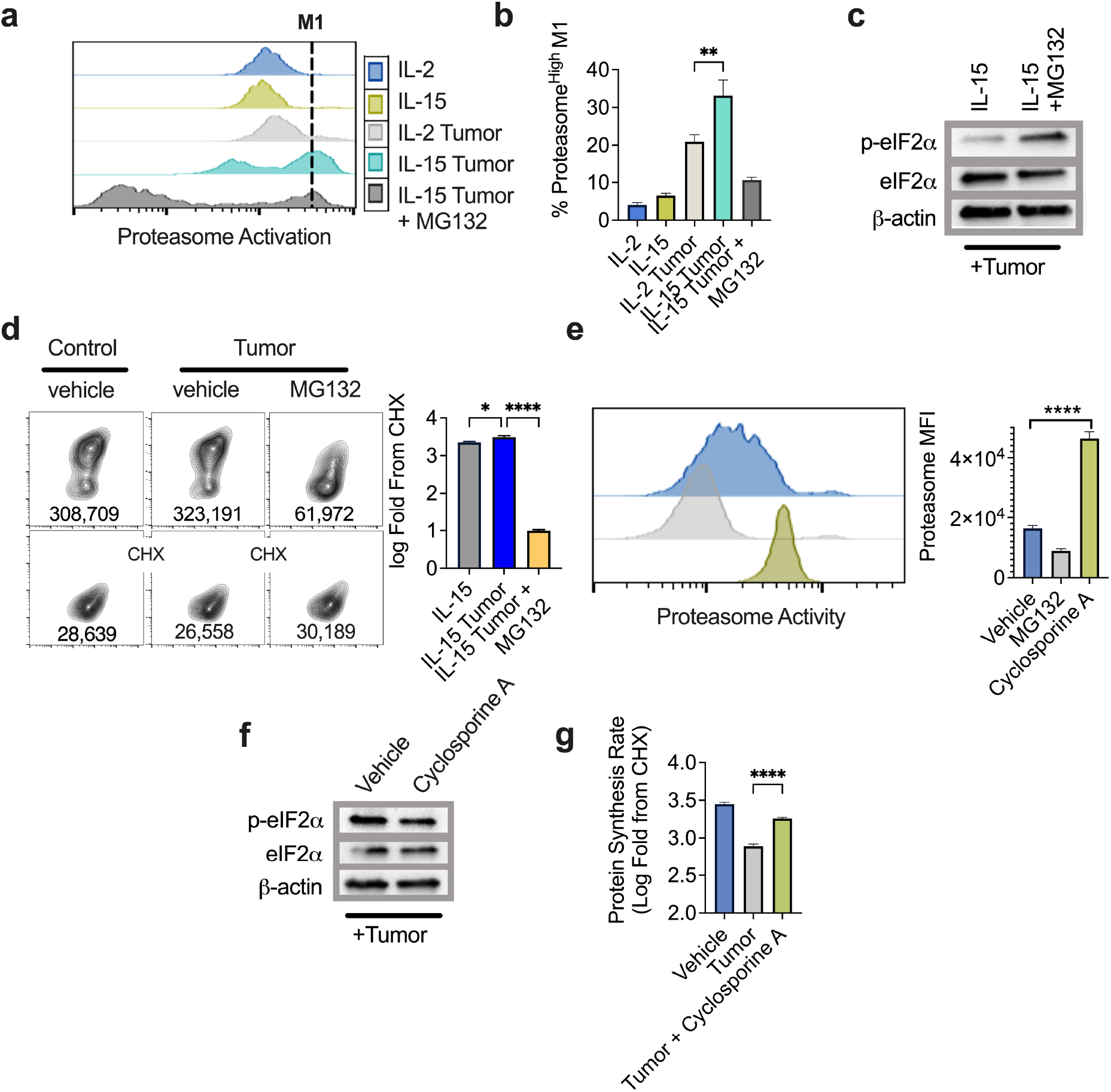
Proteasome activation represses translation attenuation. **a**) Representative FACS and **b**) quantification of the frequency of proteasome activity^high^ cells from IL-2 or IL-15 conditioned OT-1 T harvested from the tumor-T cell coculture assay. MG132 was added as an internal control 2-4 hours prior to harvest and analysis. **c**) Western blot analysis of phosphorylated or total eIF2α with β-actin loading controls and **d**) representative FACS plots and quantification of protein synthesis rates from IL-15 conditioned OT-1 T cells treated with MG132 or vehicle for 2-4 hours prior harvest from the tumor transwells. **e**) Proteasome activity of purified CD8^+^ splenocytes treated for 4 hours with cyclosporine A (2.5uM) or MG132. **f**) Western blot analysis of phosphorylated or total eIF2α with β-actin loading controls and **g**) quantification of protein synthesis rates from IL-2 effector T cells conditioned with cyclosporine A or vehicle control prior to seeding in the tumor transwell assay. Western blot or FACS plots are representative from one to three independent experiments. *p < 0.05, **p < 0.01, ***p < 0.001, ****p < 0.0001 by one way ANOVA (**b, d-e, g**). Error bars indicate the S.E.M.

### Proteasome activation represses translation attenuation

Several studies have suggested that accelerated protein degradation is a hallmark of memory T cells [22, 37], characterized by increased activation of the proteasome and enrichment for proteasome subunits. Given that optimal protein synthesis requires a delicate balance between degradation and translation in order to avoid a misfolded protein burden [38], we asked whether 26S proteasome activation could be responsible for bypassing the UPR in memory-like T cells exposed to TME stress. To assess proteasome activation in T cells we utilized a probe with fluorescent activity proportional to proteasome subunit activation [22, 39]. In tumor-seeded cocultures, IL-15-primed T cells displayed a robust increase in the frequency of proteasome activity^high^ cells (M1) relative to IL-2 effectors, which was extinguished in the presence of proteasome inhibitor MG132 (**Fig 5a-b**). Western blotting of IL-15-primed T cells revealed a substantial induction of p-eIF2α upon MG132 treatment, indicating that proteasome activation protected memory-like T cells from the UPR in tumor stress (**Fig 5c**). Indeed, proteasome inhibition in IL-15-primed T cells resulted in robust translation attenuation (**Fig 5d**), suggesting proteasome function is required to bypass the UPR and sustain synthesis.

While MG132 is a potent and selective inhibitor of proteasome function, we sought to conclusively demonstrate the importance of the proteasome function through use of pharmacologic stimulators. A recent report documented that cyclosporine A, most commonly used as an immunosuppressant, has proteasome stimulating properties which can drive memory T cell development [22]. Using the fluorescent activity probe, we validated that cyclosporine treatment increased proteasome function of naïve CD8 splenocytes (**Fig 5e**). We next addressed whether proteasome stimulation alone reduced the UPR and increased protein translation. Indeed, cyclosporine conditioning diminished p-EIF2α induction while increasing protein synthesis in T cells harvested from our tumor transwell coculture assay (**Fig 5f-g**). These findings demonstrate that proteasome activity is both necessary and sufficient to protect T cells from UPR-mediated activation of p-eIF2α, enabling sustained protein translation under TME stress.

### Proteasome function supports antitumor metabolism and immunity

Functionally, IL-15 conditioned T cells are of interest to the immunotherapy field due to their robust and prolonged levels of tumor control relative to effectors and the cytokine has been shown to enhance CAR T cell therapy [40] and response to checkpoint blockage [41]. We assessed whether access to protein degradation was a checkpoint for tumor control in memory-like T cells. Mice bearing B16F1-OVA melanomas were infused with IL-15-conditioned T cells treated with vehicle or MG132 prior to infusion and tumor growth was assessed. MG132 treatment abrogated tumor control imparted by IL-15-conditioned T cells and diminished survival benefits of the infusion (**Fig 6a-b**). In line with this, priming tumor specific effector T cells with cyclosporine A prior to adoptive transfer prolonged T cell tumor control relative to vehicle controls (**Supplemental Fig 6a**). These findings validate the importance of proteasomal function in endowing T cell mediated tumor control.

**Figure 6.**
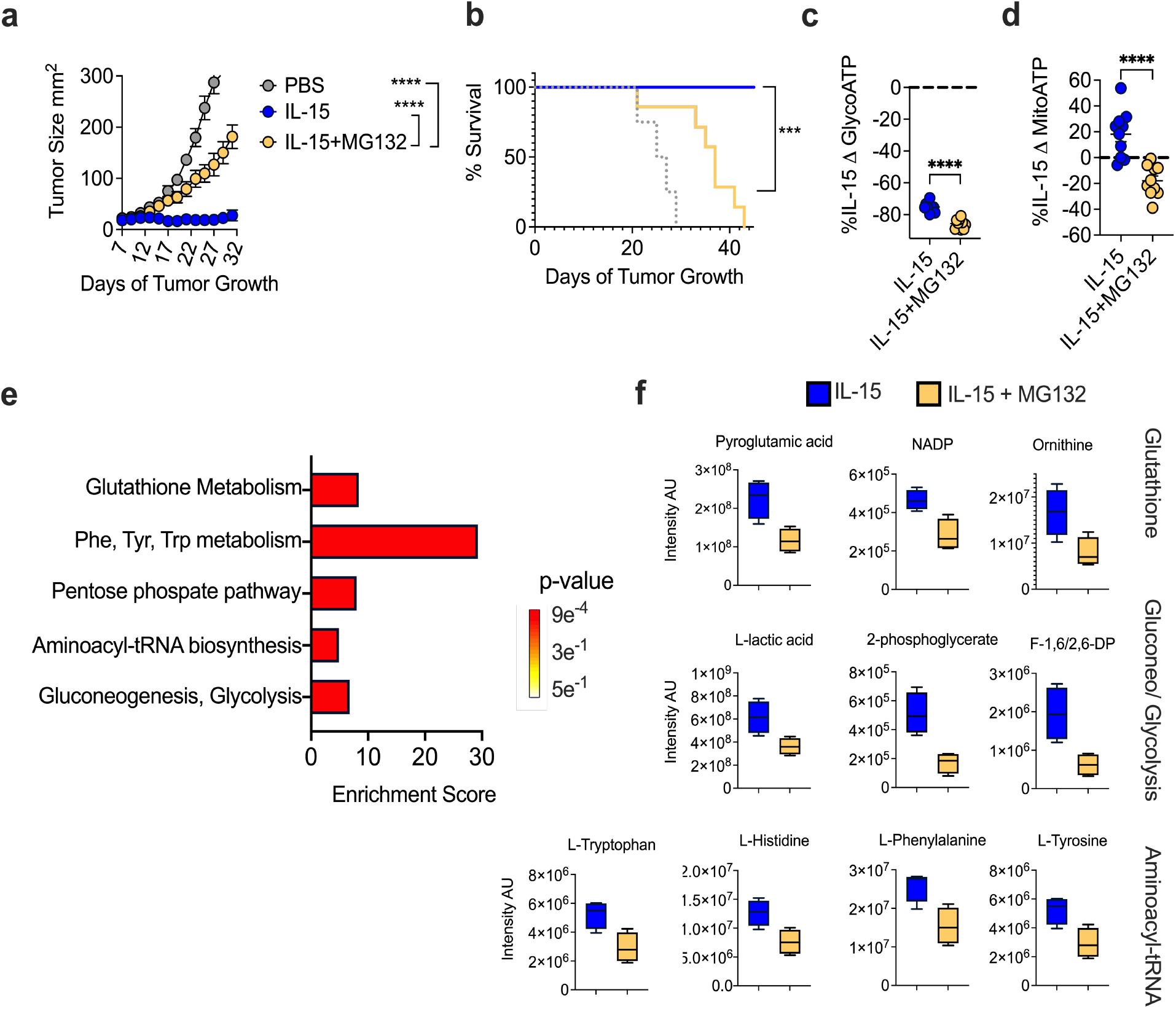
Proteasome function supports antitumor metabolism and immunity. **a**) Tumor growth rate or **b**) overall survival from adoptive transfer of day 7 expanded IL-15 conditioned OT-1 T cells treated with MG132 or vehicle control for 4 hours prior to transfer into day 7 B16-F1-OVA bearing mice (N = 4-8 mice per group). Percent change in **c**) glycolytic ATP or **d**) mitochondrial ATP of IL-15 conditioned OT1 T cells treated with MG132 relative to vehicle control from tumor transwell assay. **e**) Over representation pathway enrichment analysis of 32 metabolites upregulated in IL-15 conditioned OT-1 T cells treated with vehicle control compared to MG132 for 2-4 hours prior to harvest from the tumor transwell assay. **f**) Raw intensity values of individual metabolites upregulated in the IL-15 vehicle control cells relative to MG132 in various associated pathways. ATP rate, tumor control, and metabolomic data represent one of two independent experiments. *p < 0.05, **p < 0.01, ***p < 0.001, ****p < 0.0001 by unpaired Student’s t-test (**c-d**), linear regression (**a**), or log-rank, mantel-cox test of survival proportions (**b**). Significance cut off for determining enriched metabolites was fold change > 2 and p value < 0.05. Metabolomics data was analyzed using MetaboAnalyst 5.0 software, see methods. Error bars indicate the S.E.M

We next sought to address the mechanism by which the proteasome sustains optimal T cell function. Proteasome-mediated protein degradation supports ATP synthesis in cells experiencing nutrient stress [42]. Moreover, as shown above, IL-15 skewed T cells demonstrate a robust increase in ATP output from both glycolysis and oxidative phosphorylation compared to IL-2 effectors in the presence of tumor stress (**Fig 4**). Thus, we asked whether the proteasome was involved in metabolic remodeling of IL-15 primed cells. Proteasome inhibition with MG132 induced a significant reduction in glycolytic and mitochondrial ATP in IL-15 conditioned T cells in tumor-seeded transwells (**Fig 6c-d**). Conversely, cyclosporine A conditioned effector T cells harvested from tumor-seeded transwells demonstrated elevated levels of oxygen consumption, supporting a role for the proteasome in metabolic programming of T cells (**Supplemental Fig 6b**).

We further probed the metabolic avenues shaped by proteasome function in memory-like T cells by performing global metabolite profiling of IL-15 condition cells in tumor stress treated with vehicle of MG132. From 32 metabolites that met our criteria for significant upregulation in vehicle-treated IL-15 T cells, pathway analysis indicated upregulation of processes fundamental for memory T cell tumor control, namely gluconeogenesis and glutathione metabolism (**Fig 6e-f**) [43]. We also noted a significant enrichment in aminoacyl-tRNA biosynthesis, further validating the proteasome as an integral facet sustaining protein translation in the TME (**Fig 6e-f**). Taken together, these findings demonstrate proteasome function is necessary for optimal T cell antitumor immunity, in part through reshaping the metabolic profile of T cells in tumor stress.

## Discussion

Apart from primary translation of mRNA to amino acid chains, protein synthesis also requires successful folding of secondary, tertiary, and quaternary structures [44]. Early studies estimated that 70% of the ATP reservoirs are required to sustain translation, making it one of the most bioenergetically taxing cellular processes [45]. For T cells, antigenic stimulation activates a dynamic process highlighted by a translation rate of approximately 800,000 proteins per minute [37]. Thus, it is not challenging to recognize the substantial metabolic burden on T cells, particularly in a scenario such as the TME where competition for nutrients and global cell stressors are exceptional [5]. Despite this, very little is known about the underlying mechanisms required to support translation in the TME. We show here that blunted protein synthesis is a hallmark of tumor infiltrating CD8 T cells driven through competition for glucose leading to phosphorylation of eIF2α and downstream initiation of the UPR. These findings are a departure from previous studies of ER stress in two ways. First, we demonstrate that the TME represents a biologically relevant model to investigate the UPR in lieu of chemical insult. Second, to our knowledge, this is the first study to establish the antitumor capacity of T cells is directly related to their ability resolve ER stress. Thus, these findings implicate T cell intrinsic UPR as a previously unappreciated molecular checkpoint that dictates effective immunotherapy.

The metabolic demands of T cells are directly related to their differentiation state and predictive of their effector function [46]. Naïve T cells rely on oxidative phosphorylation, activated effector cells are highly glycolytic, and memory T cells employ *de novo* fatty acid (FA) synthesis and FA β oxidation, whereas exhausted tumor infiltrating CD8s display poor mitochondrial health associated with depressed bioenergetic output [46]. While this dependence on exogenous glucose represents a significant vulnerability to tumor infiltrating T cells, recent work has conclusively demonstrated that metabolic reprogramming cells away from glycolysis enhances cytokine production and, by extension, antitumor immunity [15, 16]. Our results here elaborate on those studies, clarifying that cytokine synthesis is likely extinguished in T cells due to glucose starvation leading to activation of the UPR branch most responsible for immediate translation attenuation. Our finding that that metabolic conditioning diminished the UPR suggests, for the first time, that T cell metabolism may be upstream of ER stress. Given the widespread interest in metabolic reprogramming as an interventional approach, the extent that UPR alleviation is necessary for efficacy with these strategies should be further elucidated. Indeed, the concepts uncovered can be extended to recent work highlighting systemic cytokine therapy rehabilitates the terminally exhausted CD8 tumor infiltrating population through reestablishing metabolic health, as connections between ER stress, the UPR, and exhausted T cells have not been made [47].

While we demonstrated metabolic reprogramming is a pathway to bypass the UPR, it is important to note this was reliant on continuous protein degradation via the 26S proteasome. In line with this, small molecule proteasome stimulators added early in the CD8 T cell activation process *in vitro* can skew cells towards a memory like phenotype [22]. Additionally, compared to naïve T cells, memory T cells isolated from human PBMC significantly upregulate translation of proteasomal subunits following stimulation [37]. Thus, our findings illustrating the importance of the proteasome for sustaining optimal antitumor immunity add to what has been established in the field. The exact mechanisms by which the proteasome supports sustained translation and tumor killing require more intricate experimentation. It is plausible that enhanced proteasome function reduces the UPR burden through cellular processes such as endoplasmic reticulum associated protein degradation [48]. Alternatively, our metabolomic data indicates a role for protein degradation in supporting metabolic pathways previously associated with optimal antitumor metabolism [43]. While the underlying mechanism requires further elucidation, our data strongly suggest proteasome stimulation as a potential method for enhancing immunotherapy outcomes.

## Methods

### Human samples

Patients undergoing surgical removal of tumors granted consent via MUSC Biorepository surgical consent forms. All study participants had not recently undergone chemo or irradiation therapy. This work was determined by MUSC Institutional Review Board to be exempt under protocol Pro00050181. Tissue samples were de-identified. Studies were conducted in accordance with the Declaration of Helsinki, International Ethical Guidelines for Biomedical Research Involving Human Subjects (CIOMS), Belmont Report, or U.S. Common Rule. Blood (8 mL) was collected in EDTA-coated tubes and PBMCs were isolated via Histopaque-1077 centrifugation. Tumor tissue was collected on ice, cut into 2-mm^3^ pieces then dissociated to single-cell suspensions using the human tumor dissociation kit and gentleMACS dissociator (Miltenyi Biotech) according to the manufacturer’s protocol. For normal human donor experiments, PBMCs and ImmunoCult Human CD3/CD28 T cell activators were obtained from Stemcell Technologies.

### Mice

OT1 (C57BL/6-Tg(TcraTcrb)1100Mjb/J), eIF2α^S51A+/-^ (B6;129-Eif2s1^tm1Rjk^/J), and C57BL/6J mice were obtained from the Jackson Laboratory. All animal experiments were approved by the Medical University of South Carolina (MUSC) Institutional Animal Care and Use Committee and the Division of Laboratory Animal Resources at MUSC maintained all mice.

### Cell cultures

B16F1 and B16F1-OVA tumor lines and OT1 T cells were maintained in RPMI supplemented with 10% FBS, 300 mg/L L-glutamine, 100 units/mL penicillin, 100 µg/mL streptomycin, 1mM sodium pyruvate, 100µM NEAA, 1mM HEPES, 55µM 2-mercaptoethanol, and 0.2% Plasmocin mycoplasma prophylactic. 0.8 mg/mL Geneticin selective antibiotic was added to media of B16F1-OVA cells for multiple passages then cells were passaged once in the Geneticin-free media prior to tumor implantation. All tumor lines were determined to be mycoplasma-free in May 2021. For OT1 T-cell activation and expansion, whole splenocytes from OT1 mice were activated with 1 µg/mL OVA 257-264 peptide and expanded for 3 days with 200 U/mL rhIL-2 (NCI). In some experiments, T cells were split on day 3 and expanded in rhIL-2, IL-15 (50ng/mL), or Cyclosporine A (2.5*μ*M) through day 7, or split to normal or low glucose (1mM) for 16 hours. Human PBMCs were activated with CD3/CD28 activators and expanded in 500U rhIL-2 (NCI), and cultures were maintained in RPMI supplemented with 10% FBS, 300 mg/L L-glutamine, 2mM GlutaMAX, 100 units/mL penicillin, 100 µg/mL streptomycin, 50 µg/mL gentamycin, 25mM HEPES, 55µM 2-mercaptoethanol, and 0.2% Plasmocin mycoplasma prophylactic.

### Tumor models

For endogenous TIL analysis, B16F1 were established by injecting 2.5×10^5^ cells subcutaneously into the right flank of female C57BL/6 mice. Tumor-draining lymph nodes, spleens, and tumors were harvested after 6 days of tumor growth for measurement of protein synthesis. For adoptive cellular therapy experiments, B16F1-OVA melanomas were established subcutaneously by injecting 2.5×10^5^ cells into the right flank of female C57BL/6 mice and tumor-bearing hosts were irradiated with 5 Gy 24 hours prior to T-cell transfer. After 7 days of tumor growth, 5×10^5^ OT1 T cells conditioned with IL2, IL15, vehicle, or Cyclosporin A were infused in 100 µL phosphate-buffered saline via tail vein into mice. For proteasome inhibition, vehicle or MG132 (5mM) was added to T cells 4 hours prior to infusion. Tumor growth was measured every other day with calipers, and survival was monitored with an experimental endpoint of tumor growth ≥ 300 mm^2^

### Tumor-T cell transwell assays and glucose measurement

4×10^5^ or 1×10^5^ B16F1 tumor cells were seeded into 6 or 12 well companion plates, respectively, 24 hours prior to addition of OT1 T cells or PBMCs in transwells inserts in complete T-cell media supplemented with 200 U rhIL-2 (NCI) or IL15 (50ng/mL) and harvested 36 hours later. Mouse T cells were added to transwells at peptide activation, and in other instances added after 3 days of expansion in 200 U/mL rhIL-2 (NCI). Human T cells were added to transwells at soluble CD3/28 activation, and in other instances added after 1 week of expansion. For metabolic remodeling, high glucose (25mM), 2DG (1mM, chronic), IL15 (50nM), or Cyclosporine A (2.5*μ*M) were introduced at the time of T cell addition to transwells. For acute treatment, 2DG (1mM, acute), or MG132 (5*μ*M) were added 2 and 4 hours prior to bioenergetic analysis, respectively. Glucose concentration in transwell assay media with increasing tumor density was determined using the B61000 Blood Glucose System on the day of transwell assay harvest. L-amino acid concentrations in media were determined using the L-amino Acid Assay Kit (Abcam) according to manufacturer’s protocol.

### Immunoblotting

T cells were lysed in RIPA Buffer (Sigma) supplemented with Protease Inhibitor Cocktail (Cell Signaling Technology) and Phosphatase Inhibitors I and II (Sigma). Protein concentrations were normalized using Pierce BCA Kit (Thermo Fisher Scientific) and loaded to 4%–10% agarose gels (Bio-Rad). p-eIF2α, eIF2*α, β*-actin, and HRP-linked anti-rabbit and mouse secondaries were obtained from Cell Signaling Technology. Phospho protein was developed with Pierce ECL Plus Western Blotting Substrate (Thermo Fisher Scientific).

### Protein synthesis and flow cytometry

Homopropargylglycine (HPG) protein synthesis was measured using the Click-iT HPG Alexa Fluor 488 Protein Synthesis Assay Kit from Thermo Fisher Scientific. Cells were incubated in methionine-free media and controls were treated with cycloheximide. Cells were stained extracellularly using CD8-Alexa Fluor 647, CD8-eFluor 710, CD45.2-Alexa Fluor 647, and PD-1-eFluor 710 or isotype control and subsequently incubated in 50µM L-HPG with added Live-or-Dye Fixable Viability Stain. Cells were fixed using Fixation Buffer (Biolegend) and permeabilized with Triton X-100 solution followed by staining with Alexa Fluor 488 Azide to label active protein synthesis. OPP protein synthesis was measured using the O-Propargyl-puromycin (OPP) Protein Synthesis Assay Kit (Cayman Chemical). T cells were incubated in complete T-cell media, and control cells were treated with CHX. Cells were incubated in cell-permeable OPP, and Live-or-Dye Fixable Viability Stain was added. Cells were fixed using formaldehyde and subsequently stained with 5 FAM-Azide to label translating polypeptide chains. Cells were stained extracellularly using CD8-Alexa Fluor 647. Samples were run directly on a BD Accuri C6 flow cytometer and analysis was performed with FlowJo software (BD Biosciences). For proteasome activity analysis, Proteasome Activity Probe (R&D Systems) was stained on cells of interest at 2.5*μ*M for 2 hours at 37 C in PBS. Samples were run directly on a BD Accuri C6 flow cytometer and analysis was performed with FlowJo software.

### Cytokine synthesis

For cytokine restimulated human and mouse T cells, in some instances cells grown in normal media or tumor supernatant were treated with cycloheximide for 30 minutes prior to being restimulated for 4 hours with Cell Stimulation Cocktail and GolgiPlug Brefeldin A (eBioscience). Cells were stained extracellularly then IFN*γ* TNF-α intracellular FACS staining was performed using the Foxp3/Transcription Factor Fixation/Permeabilization Concentrate and Diluent kit and Permeabilization Buffer (eBioscience). Samples were run on a BD Accuri C6 flow cytometer and analysis was performed with FlowJo software.

### Seahorse bioanalysis

Seahorse XF Real-Time ATP Rate assays were performed using the Seahorse XFe96 analyzer. 96-well plates were coated with CellTak (Corning), washed, and air dried. T cells were plated in Seahorse XF DMEM Medium (Agilent) supplemented with 1% fetal bovine serum (FBS) and centrifuged for adherence. Next, 1µM oligomycin and 2µM rotenone/1µM antimycin A were injected sequentially, and the oxygen consumption rate (OCR) and extracellular acidification rate (ECAR) were measured. The mitochondrial ATP production rate was quantified based on the decrease in the OCR. The glycolytic ATP production rate was calculated as the increase in the ECAR combined with total proton efflux fate (PER). Seahorse XF Cell Mito Stress Test assays were performed as above and 1µM oligomycin, 1.5uM FCCP, and 2µM rotenone/1µM antimycin A were injected sequentially, and the oxygen consumption rate (OCR) was measured. Spare respiratory capacity was calculated as the difference between the basal and maximal OCR readings after addition of FCCP.

### RNA analysis and UPR arrays

RNA was isolated with RNeasy Mini Kit (Qiagen, 74104) and concentration was measured using the SpectraDrop Micro-Volume Microplate (Molecular Devices). Single-strand cDNA was made with 500 ng RNA using the High Capacity RNA-to-cDNA Kit (Applied Biosystems, 4387406, Thermo Fisher Scientific). TaqMan Gene Expression Assays (Applied Biosystems, Thermo Fisher Scientific) were used to perform single-gene real-time PCR and TaqMan Gene Expression Array Plates (Thermo Fisher Scientific) were used to perform Unfolded Protein Response (UPR) arrays. All experiments were performed using the StepOnePlus Real-Time PCR System (Applied Biosystems, Thermo Fisher Scientific). Gene expression was normalized to Gapdh for single gene assays and Gapdh, Gusb, β-Actin, and β2M for UPR arrays and results were calculated using ΔΔCt method.

### Metabolomics

#### Sample Preparation

Cells were harvested and washed with sodium chloride solution then resuspended in 80% methanol for multiple freeze-thaw cycles at −80°C and stored at −80°C overnight to precipitate the hydrophilic metabolites. Samples were centrifuged at 20,000xg and methanol was extracted for metabolite analysis. Remaining protein pellets were dissolved with 8M urea and total protein was quantified using BCA assay. Total protein amount was used for equivalent loading for High Performance Liquid Chromatography and High-Resolution Mass Spectrometry and Tandem Mass Spectrometry (HPLC-MS/MS) analysis.

#### Data Acquisition

Samples were dried with a SpeedVac then 50% acetonitrile was added for reconstitution followed by overtaxing for 30 sec. Sample solutions were centrifuged and supernatant was collected and analyzed by High-Performance Liquid Chromatography and High-Resolution Mass Spectrometry and Tandem Mass Spectrometry (HPLC-MS/MS). The system consists of a Thermo Q-Exactive in line with an electrospray source and an Ultimate3000 (Thermo) series HPLC consisting of a binary pump, degasser, and auto-sampler outfitted with a Xbridge Amide column (Waters; dimensions of 2.3 mm × 100 mm and a 3.5 µm particle size). The mobile phase A contained 95% (vol/vol) water, 5% (vol/vol) acetonitrile, 10 mM ammonium hydroxide, 10 mM ammonium acetate, pH = 9.0; B with 100% Acetonitrile. For gradient: 0 min, 15% A; 2.5 min, 30% A; 7 min, 43% A; 16 min, 62% A; 16.1-18 min, 75% A; 18-25 min, 15% A with flow rate of 150 μL/min. A capillary of the ESI source was set to 275 °C, with sheath gas at 35 arbitrary units, auxiliary gas at 5 arbitrary units and the spray voltage at 4.0 kV. In positive/negative polarity switching mode, an *m*/*z* scan range from 60 to 900 was chosen and MS1 data was collected at a resolution of 70,000. The automatic gain control (AGC) target was set at 1 × 10^6^ and the maximum injection time were 200 ms. The top 5 precursor ions were subsequently fragmented, in a data-dependent manner, using the higher energy collisional dissociation (HCD) cell set to 30% normalized collision energy in MS2 at a resolution power of 17,500. Besides matching m/z, metabolites were identified by matching either retention time with analytical standards and/or MS2 fragmentation pattern. Metabolite acquisition and identification was carried out by Xcalibur 4.1 software and Tracefinder 4.1 software, respectively.

#### Statistical Analysis

Post identification, samples were normalized by taking the peak area under the curve for each metabolite per sample and dividing by the quotient of the total ion count (TIC) per sample over the lowest TIC in the batch. Subsequent transformation of normalized data was carried out with auto scaling to account for heteroscedasticity [49]. Metabolites that were below detection in all samples were removed from analysis; missing values were imputed with 1/5 of the minimum positive value of their corresponding variable. Differential metabolite expression between groups of interest were identified through a combination of fold change >2 and raw p-value < 0.1 and visualized using a clustered heatmap. Overrepresentation enrichment analysis using the KEGG database was performed on metabolites meeting this criterion to identify biological processes associated with differential expression [50]. All statistical analysis was performed using the MetaboAnalyst 5.0 web server [51].

### Statistical Analysis

GraphPad Prism was used to calculate p values with one-way analysis of variation (ANOVA) with Dunnett’s multiple comparison test, unpaired two-sided Student’s *t-*test or paired Student’s *t-*test as indicated in the figure legend. For all statistical analyses, the assumption of Gaussian distribution was validated using the *F* test to compare variance or the Brown-Forsythe test for Student’s *t* Test or one-way ANOVA, respectively. Tumor growth curves and survival were calculated using linear regression and Log-Rank test, respectively. All data are presented as mean +/- Standard Error of the Mean (S.E.M.) unless otherwise denoted. Values with an α < 0.05 were considered significant. Significant p values were denoted as: * p < 0.05, ** p < 0.01, *** p < 0.001, **** p < 0.0001.

## Acknowledgements

We are thankful to Dr. Zihai Li for continued mentorship over the course of this work. We acknowledge the supported in part by the Biostatistics Shared Resource, Hollings Cancer Center, Medical University of South Carolina (P30 CA138313).

## Notes

Financial Support: R01CA244361-01A1, R01CA248359-01 (JET), T32 5T32AI132164-04 (BPR), T32 DE01755 (MDT), T32 CA 193201 (AMA)

### Competing Interest Statement

The authors have declared no competing interest.

