## Supplemental Information for "Stress-Mediated Attenuation of Translation Undermines T Cell Tumor Control"

**Running Title**: Translation attenuation impedes antitumor immunity

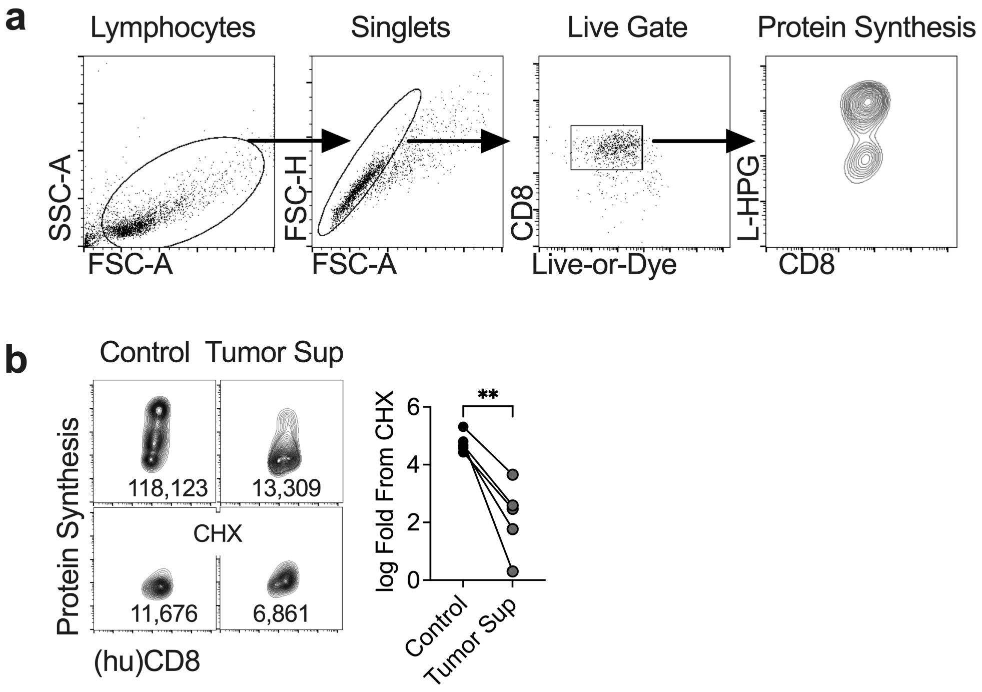


**Supplemental Figure 1. Tumor supernatant blunts T cell protein synthesis**
**a**) Gating strategy for quantification of protein synthesis in live CD8 T cells harvested from the tumor transwell assay. **b**) Representative FACS plots and quantification of protein synthesis from human CD8 PBMC (n=5 donors) activated with CD3/CD28 in the presence or absence of supernatant from freshly isolated B16F1 melanoma tumors. ** $p < 0.01$, two tailed paired Student's *t* test.

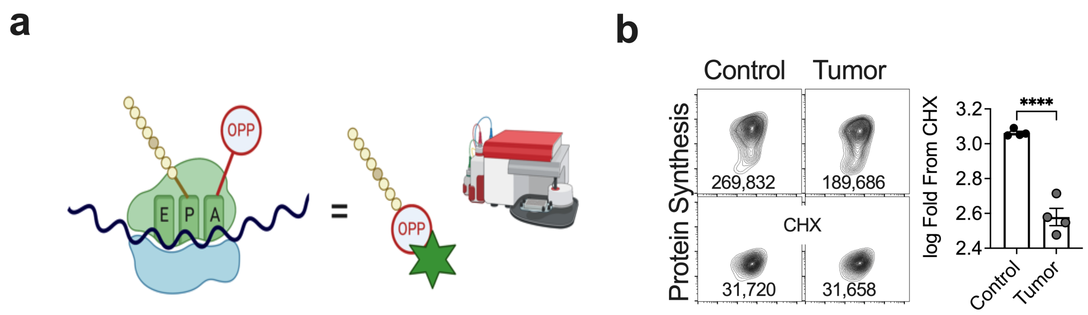


**Supplemental Figure 2. Validation of tumor mediated reduction in protein synthesis**
**a**) Schematic representation of the O-Propargyl-puromycin (OPP) assay used to measure protein synthesis. **b**) Representative FACS plots and quantification of OPP protein synthesis from OT-1 T cells harvested from the tumor transwell assay. **** $p < 0.0001$, two tailed Student's *t* test. Error bars represent S.E.M.

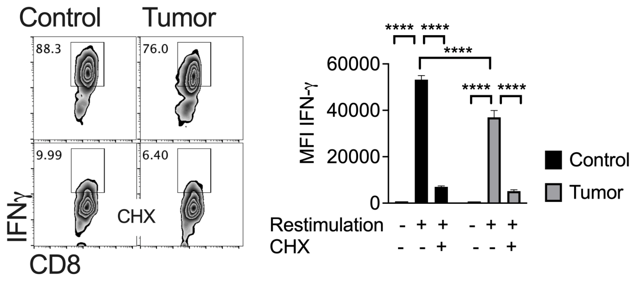


**Supplemental Figure 3**. **Reduced IFNγ production driven by TME pressure**
Representative FACS plots and quantification of IFNγ production from OT-1 T cells harvested from the tumor transwell assay followed by stimulation with PMA/Ionomycin ± pretreatment with protein synthesis inhibitor cycloheximide (CHX). **** $p < 0.0001$ by one-way ANOVA. Error bars represent S.E.M.

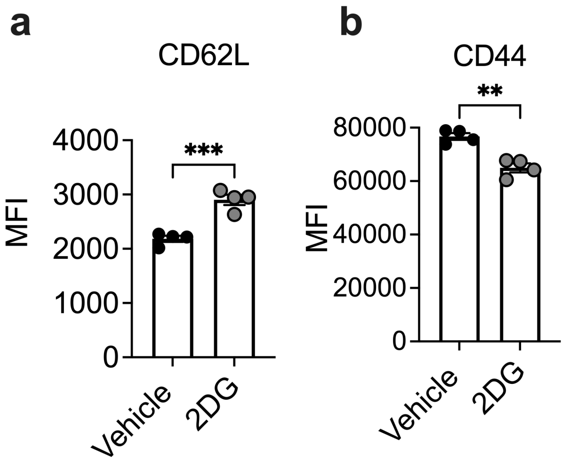


**Supplemental Figure 4. Phenotypic analysis of 2DG conditioned CD8 T cells**
Mean fluorescent intensity of **a**) CD62L and **b**) CD44 on OT-1 T cells treated with vehicle or 2DG for 36 hours. ** $p < 0.01$, *** $p < 0.001$ by two tailed Student's *t* test. Error bars represent S.E.M.

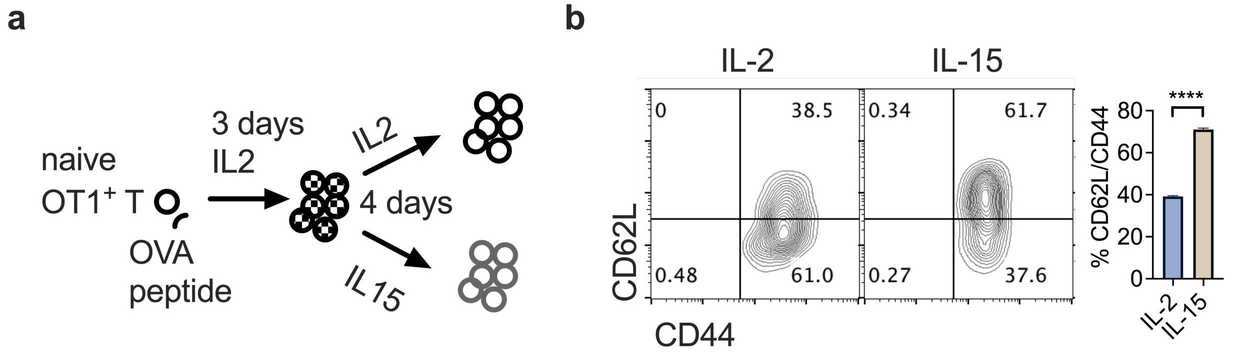


**Supplemental Figure 5. IL-15 conditioning enhances memory-like T cell phenotype**
**a**) Graphical representation of the memory or effector like cytokine conditioning scheme. **b)** Representative FACS plots and quantification of CD44 and CD62L expression on IL-2 or IL-15 conditioned OT-1 T cells as demonstrated in (**a**). **** $p < 0.0001$ by two tailed Student's *t* test. Error bars represent S.E.M.

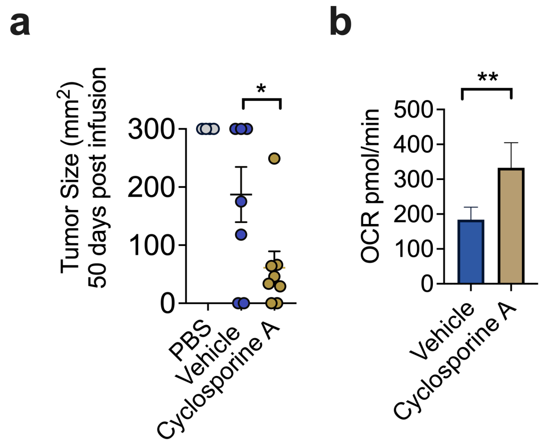


**Supplemental Figure 6. Proteasome stimulation enhances antitumor immunity**
**a**) B16-F1-OVA tumor sizes in mice 50 days post infusion of day 7 activated OT-1 T cells conditioned with vehicle or 2.5μM of Cyclosporine A prior to infusion. Day 7 tumor bearing mice were sublethally irradiated 24 hours prior to adoptive transfer. **b**) Seahorse bioanalysis to quantify spare respiratory capacity of OT-1 T cells treated with vehicle or 2.5μM Cyclosporine A for 4 days prior. * $p < 0.05$, ** $p < 0.01$, by one-way ANOVA (**a**) or two tailed Student's t Test (**b**). Error bars represent S.E.M.
